# Whole genome linkage disequilibrium and effective population size in a coho salmon (*Oncorhynchus kisutch*) breeding population

**DOI:** 10.1101/335018

**Authors:** Agustín Barría, Kris A. Christensen, Grazyella Yoshida, Ana Jedlicki, Jean P. Lhorente, William S. Davidson, José M. Yáñez

**Affiliations:** Facultad de Ciencias Veterinarias y Pecuarias, Universidad de Chile, La Pintana, Santiago, Chile; Department of Molecular Biology and Biochemistry, Simon Fraser University, Burnaby, BC, Canada; Animal Science Department, Universidade Estadual Paulista “Júlio de Mesquita Filho” (UNESP), Faculdade de Ciências Agrárias e Veterinárias (FCAV), Campus Jaboticabal, Via de Acesso Prof. Paulo Donato Castellane, Jaboticabal, Brazil; Aquainnovo S.A., Puerto Montt, Chile.; Núcleo Milenio INVASAL, Concepción, Chile

**Keywords:** Linkage disequilibrium, *Oncorhynchus kisutch*, Selective breeding, GWAS, Effective population size

## Abstract

The estimation of linkage disequilibrium between molecular markers within a population is critical when establishing the minimum number of markers required for association studies, genomic selection and for inferring historical events influencing different populations. This work aimed to evaluate the extent and decay of linkage disequilibrium in a coho salmon breeding population using ddRAD genomic markers.

Linkage disequilibrium was estimated between a total of 7,505 SNPs found in 62 individuals (33 dams and 29 sires) from the breeding population. The makers encompass all 30 coho salmon chromosomes and comprise 1,655.19 Mb of the genome. The average density of markers per chromosome ranged from 3.45 to 6.11 per 1 Mbp. The minor allele frequency averaged 0.20 (with a range from 0.08 to 0.50). The overall average linkage disequilibrium among SNPs pairs measured as r^2^ was 0.054. The Average r^2^ value decreased with increasing physical distance, with values ranging from 0.37 to 0.054 at distances lower than 1 kb and up to 10 Mb, respectively. An r^2^ threshold of 0.1 was reached at distance of approximately 1.3 Mb. Chromosomes Okis05, Okis15 and Okis28 showed high levels of linkage disequilibrium (> 0.20 at distances lower than 1 Mb). Average r^2^ values were lower than 0.1 for all chromosomes at distances greater than 4 Mb. Linkage disequilibrium values suggest that whole genome association and selection studies could be performed using about 75,000 SNPs in aquaculture populations (depending on the trait under investigation). From the identified SNPs, an effective population size of 100 was estimated for the population 10 generation ago, and 1,000, for 139 generations ago.

Based on the extent of r^2^ decay, we suggest that at least 75,000 SNPs would be necessary for an association mapping study. Over 100,000 SNPs would be necessary for a high power study, in the current coho salmon population.

## INTRODUCTION

Coho salmon (*Oncorhynchus kisutch*) is one of the six Pacific salmon species found in North American and Asian watersheds (Groot and Margolis, 1991). This species was introduced into Chilean streams during the 1920s promoted by the Chilean Institute of Fisheries Department. Cultivation of coho salmon began in Chile at the end of the 1970’s, when Chile imported almost 500,000 eggs from the Kitimat river (British Columbia) and Oregon, becoming the genetic basis of the broodstocks in Chile (Neira et al., 2014). Twenty years later, the production of the first eggs for commercial use were produced in Chile (SalmonChile, 2007). Currently, Chile is the main producer of farmed coho salmon, with the production of nearly 160,000 tons in 2014 (FAO, 2016). This represents more than 90% of the global farmed coho production (Canada and Japan are the other major coho salmon producers) (FAO, 2016). The temperature and the quality of the Chilean freshwater environments have reduced the coho reproductive cycle to only two years (Estay et al., 1997). To date, numerous genetic programs have been developed for coho salmon in Chile. These programs are mainly focused on growth, disease resistance, and flesh color (Neira et al., 2014).

With the eruption of next generation sequencing (NGS) technologies, it has become possible to perform artificial selection through the use of genomic estimated breeding values (GEBVs). By using dense molecular markers from the whole genome, genomic selection (GS) can be used in broodstock enhancement (Bennewitz et al., 2009). This methodology makes it possible to estimate GEBVs with high accuracy, even with animals without recorded phenotypes (Meuwissen et al., 2001), which has improved the accuracy of selection in salmonid species (Bangera et al., 2017; Barría et al., 2018; Correa et al., 2017; Ødegård et al., 2014; Tsai et al., 2016; Yoshida et al., 2018). Genome wide association studies (GWAs) and GS, exploit linkage disequilibrium (LD) between molecular markers. The amount of LD between loci is important in GWAs, as the extent of LD indicates the necessary number of SNPs to assure that causative mutations are in LD with genetic markers (Flint-Garcia et al., 2003). GWAs are key for mapping traits with commercial interest to causative mutations in the genome. For GS, LD is related to the likelihood of successfully tagging the SNP effect in genomic breeding value prediction (Kemper and Goddard, 2012).

LD maps allow researchers to explore the genetic basis of traits influencing productivity. Through the comparison of the extent and pattern of LD in these LD maps, it is possible to elucidate the diversity among breeds with different phenotypic attributes, and even identify genomic regions subject to different selective pressures (López et al., 2015; McKay et al., 2007). The most common LD measurements are r^2^ and |D’|, both ranging from 0 to 1. When |D’| < 1, it indicates the occurrence of historical recombination between loci, while |D’| = 1 indicates no recombination. The r^2^ statistic represents the correlation between molecular markers. This latter parameter is preferred over |D’| because |D’| tends to be overestimated in small samples size and when low-frequencies alleles are used (Teare et al., 2002). Moreover, in association studies, r^2^ is preferred due to the inverse relationship between its value and the sample size needed to detect a significant association between a causative variant and molecular markers (Wall and Pritchard, 2003).

Despite the many GWAs and GS analyses performed with Atlantic salmon (Bangera et al., 2017; Correa et al., 2017; Gutierrez et al., 2015; Tsai et al., 2015, 2016), rainbow trout (Vallejo et al., 2016, 2017) and Coho salmon (Barría et al., 2018), none of them have evaluated the LD in the studied populations. Further, most of the linkage disequilibrium studies have been focused on the extent and decay pattern of LD in livestock species, such as dairy (Bohmanova et al., 2010; Sargolzaei et al., 2008) and beef cattle (Makina et al., 2015; McKay et al., 2007), plants (Delourme et al., 2013; Porto-Neto et al., 2014), and pigs (Saura et al., 2015). Recently, LD has been evaluated in farmed rainbow trout (*Oncorhynchus mykiss*) (Rexroad and Vallejo, 2009) and in Atlantic salmon (Kijas et al., 2017).

The first step to calculate the number of molecular markers necessary for genomic selection and QTL mapping is to estimate the extent and decline of LD within a population. To date, there have been no studies aimed to characterize the levels and extent of LD in coho salmon. The current work aimed to evaluate the extent of linkage disequilibrium, at the genomic and chromosome level, on a breeding coho salmon population using double digest restriction associated DNA (ddRAD) molecular markers.

## MATERIAL AND METHODS

### Populations and samples

The coho salmon samples were obtained from a breeding population belonging to a genetic improvement program established in 1997. Using *best linear unbiased prediction* (BLUP), harvest weight had been selected over eight generations in this population. For LD estimations a total of 63 animals (33 sires and 30 dams), corresponding to the parents of 33 families from a 2012-spawning year, were selected. These individuals were not related to each other and belonged to the broodstock of the breeding nucleus. For specific details about reproductive management, mating design, rearing conditions, effective size, inbreeding and breeding objectives of the genetic program for this population see (Dufflocq et al., 2016; Yáñez et al., 2014, 2016). Experimental challenge against *P. salmonis* was approved by the Animal Bioethics Committee from Universidad de Chile (N°08-2015).

### Genotyping

Total DNA was extracted from the fin clip of 63 individuals. Sequencing library preparation was performed using a ddRAD methodology, and followed the protocol described in Peterson et al., (2012). Briefly, the extracted and normalized DNA was digested with the restriction enzymes (RE) *SbfI* and *MseI,* followed by adapter ligation and PCR amplification with primers complementary to the adapters. After PCR, individual sample libraries were pooled and concentrated. Size selection was performed on the pooled libraries (from 0.75 and 1.5 kb and between 1.8 and 2.5 kb). Sequencing was performed on a HiSeq2500 using 150 bp SE.

For the SNP identification step, raw sequences were analyzed with STACKS v. 1.41 (Catchen et al., 2011, 2013). Reads were trimmed to 134 bp using *process_radtags*. After this quality filter step, the sequences were aligned to the coho salmon reference genome (assembly accession = GCA_002021735.1) using BWA mem (Li and Durbin, 2009). For loci identification, a minimum coverage depth of three (-m 3) was set in *pstacks*. To build the genetic marker catalog, reads from all genotyped fish were considered and processed with *cstacks*. The *sstacks* program was then used to match called individual loci against the constructed catalog, and genotypes were called with the *populations* program. Loci were considered only if present in at least 75% of the individuals. Prior to LD analyses, a quality control (QC) step was performed through the GenABEL library (Aulchenko et al., 2007) implemented in R (R Core Team, 2016). The following parameters were used to exclude low-confidence SNPs: Hardy-Weinberg Equilibrium (HWE) p < 1e-6, Minor Allele Frequency (MAF) ≤ 0.05 and genotyping call rate < 0.80. Fish with genotyping call rates < 0.70 were excluded from further analyses. For more details about DNA extraction, library preparation, data filter and SNP quality, please see (Barría et al., 2018). With the availability of the coho salmon reference genome, physical distances were calculated during SNP identification. Raw sequences were deposited at the NCBI Sequence Read Archive (SRA). Bioproject ID PRJNA471180, temporary submission ID SUB4039075.

### LD estimation

The LD between each pair of genetic markers was estimated using Pearson’s squared correlation coefficient (r^2^) statistic which is less sensitive to allelic frequencies (Ardlie et al., 2002) and more suitable for biallelic markers (Zhao et al., 2005). The r^2^ values were estimated with Plink v1.09 (Purcell et al., 2007). Genotypes were coded as 0, 1 and 2 relative to the number of non-reference alleles. The parameter –inter-chr, in conjunction with a ld-window-r2 set to zero, was used to obtain correlations between all the pairs of SNPs on each chromosome independently of their r^2^ value.

LD decay curves were calculated for SNP pairs separated by an inter-marker distance between 0 and 10 Mb, and per chromosome according to the distance between all markers on that chromosome.

The historical effective population size was estimated using SNeP (Barbato et al., 2015). Using the estimated LD values, a historical population size estimation was calculated with the following equation: N_t_ = 1/(4*f*(c)) (1/r^2^ – 1). Where *f*(c) refers to c[(1 - c/2)/(1 - 2)^2^] (Sved, 1971), and *c* corresponds to the linkage distance inferred from the physical distance between SNPs, assuming that 1 Mb ∼ 1cM and that N_t_ is the effective population size estimated at t = 1/2c generations ago.

## RESULTS

### SNPs identification

A total of 9,376 SNPs was identified in 62 coho salmon individuals (one individual was dropped). Of these, 7,505 were on chromosomes. The rest of the markers (1,871) were excluded because they mapped to unplaced scaffolds (i.e. the scaffolds were not on chromosomes). Markers tented to have low MAF values between 0.05 and 0.1. The MAF distribution of the retained SNPs was nearly uniform along the 30 chromosomes, with an average of 0.20 ± 0.13 (mean ± standard deviation), and a minimum and maximum value of 0.18 and 0.29, respectively (**Table 1**). The average decreased to 0.19 ± 0.12 when all 9,376 SNPs were included.

**Table 1.** Summary statistics for the evaluated SNPs and linkage disequilibrium values along coho salmon chromosomes.

| Okis | Length (Mb) | Number of SNPs | SNP Density (Mb) | Mean ($r^2$) | Median ($r^2$) | SD ($r^2$) | MAF |
| --- | --- | --- | --- | --- | --- | --- | --- |
| 01 | 66.78 | 299 | 4.47 | 0.061 | 0.023 | 0.12 | 0.20 |
| 02 | 73.79 | 299 | 4.05 | 0.050 | 0.020 | 0.10 | 0.19 |
| 03 | 68.18 | 288 | 4.22 | 0.056 | 0.021 | 0.11 | 0.20 |
| 04 | 79.31 | 404 | 5.09 | 0.048 | 0.019 | 0.09 | 0.18 |
| 05 | 70.56 | 253 | 3.59 | 0.057 | 0.020 | 0.11 | 0.18 |
| 06 | 76.16 | 314 | 4.12 | 0.048 | 0.016 | 0.10 | 0.20 |
| 07 | 50.17 | 232 | 4.62 | 0.053 | 0.020 | 0.10 | 0.20 |
| 08 | 66.26 | 283 | 4.27 | 0.061 | 0.023 | 0.12 | 0.20 |
| 09 | 39.12 | 161 | 4.12 | 0.052 | 0.019 | 0.11 | 0.18 |
| 10 | 64.07 | 324 | 5.06 | 0.048 | 0.018 | 0.10 | 0.20 |
| 11 | 78.70 | 362 | 4.60 | 0.065 | 0.025 | 0.12 | 0.21 |
| 12 | 50.46 | 281 | 5.57 | 0.051 | 0.021 | 0.10 | 0.19 |
| 13 | 66.72 | 333 | 5.00 | 0.045 | 0.017 | 0.10 | 0.19 |
| 14 | 69.58 | 242 | 3.48 | 0.055 | 0.021 | 0.10 | 0.20 |
| 15 | 64.02 | 287 | 4.48 | 0.051 | 0.020 | 0.10 | 0.20 |
| 16 | 32.35 | 145 | 4.48 | 0.056 | 0.019 | 0.12 | 0.20 |
| 17 | 75.25 | 344 | 4.57 | 0.054 | 0.020 | 0.11 | 0.21 |
| 18 | 64.85 | 303 | 4.67 | 0.051 | 0.019 | 0.10 | 0.20 |
| 19 | 52.74 | 226 | 4.29 | 0.060 | 0.023 | 0.12 | 0.19 |
| 20 | 40.39 | 226 | 5.60 | 0.043 | 0.015 | 0.10 | 0.18 |
| 21 | 34.84 | 165 | 4.74 | 0.051 | 0.017 | 0.11 | 0.18 |
| 22 | 55.31 | 228 | 4.12 | 0.056 | 0.022 | 0.10 | 0.20 |
| 23 | 41.04 | 196 | 4.78 | 0.051 | 0.019 | 0.11 | 0.20 |
| 24 | 38.72 | 172 | 4.44 | 0.055 | 0.020 | 0.11 | 0.19 |
| 25 | 32.99 | 201 | 6.10 | 0.055 | 0.023 | 0.11 | 0.18 |
| 26 | 43.33 | 181 | 4.18 | 0.060 | 0.022 | 0.12 | 0.20 |
| 27 | 37.31 | 228 | 6.11 | 0.061 | 0.024 | 0.11 | 0.20 |
| 28 | 46.42 | 190 | 4.10 | 0.062 | 0.020 | 0.13 | 0.19 |
| 29 | 36.55 | 182 | 4.98 | 0.048 | 0.018 | 0.10 | 0.18 |
| 30 | 39.22 | 156 | 3.98 | 0.062 | 0.021 | 0.13 | 0.21 |
| Mean | 55.17 | 250 | 4.60 | 0.054 | 0.020 | 0.11 | 0.20 |

SNP: Single-Nucleotide Polymorphism; MAF: Minor Allele Frequency; SD: Standard deviation

### Estimation of LD

**Table 1** summarizes the mean, median and standard deviation of r^2^ values for each coho salmon chromosome. All of the 7,505 SNPs placed onto chromosomes and which passed quality control were included in this analysis. These markers encompassed 1,655.19 Mb of the genome. The molecular marker density per chromosome per Mb, ranged from 3.48 to 6.11 with a mean of 4.60. In general, SNPs were uniformly distributed along the 30 chromosomes. The number of SNPs on each chromosome ranged from 145 on Okis16 to 404 on Okis04, which is in agreement with Okis16 and Okis04 being the shortest and longest chromosome, respectively. The overall mean linkage disequilibrium (measure as r^2^) among SNP pairs was 0.054 ± 0.11. The global median was lower at 0.020. Low average LD among adjacent SNPs along the 30 chromosomes was observed in the current population, with values ranging from 0.043 to 0.065 (**Table 1**).

In order to estimate the decay of linkage disequilibrium as a function of physical distance, SNP pairs were sorted into bins based on their distance, and mean values of r^2^ were calculated for each bin. As observed in other species (Kijas et al., 2017; Lu et al., 2012; Vos et al., 2017), LD declines smoothly as the physical distance increases between markers. **Figure 1** shows a scatter plot of the decline in r^2^ among linked SNP pairs as distance increases. A maximum average LD of 0.37 was estimated for SNPs less than 1 kb apart. This value declines quickly by more than half at marker distances up to 0.1 Mb, with a value of 0.12. From 1 Mb to 10 Mb LD ranges from 0.103 to 0.051. The latter value represents the lowest average LD estimated in the current data set. The r^2^ estimation drops below 0.1 at a distance of ∼ 1.3 Mb, suggesting that linkage equilibrium was reached (Vos et al., 2017). **Figure 2** compares the mean LD at different distance bins for each chromosome. High variation across chromosomes was observed at lower distance bins. Higher levels of LD (> 0.20) were estimated for Okis05, Okis15 and Okis28. Average r^2^ values < 0.10 were estimated for all chromosomes at distances greater than 4 Mb.

**Figure 1.**
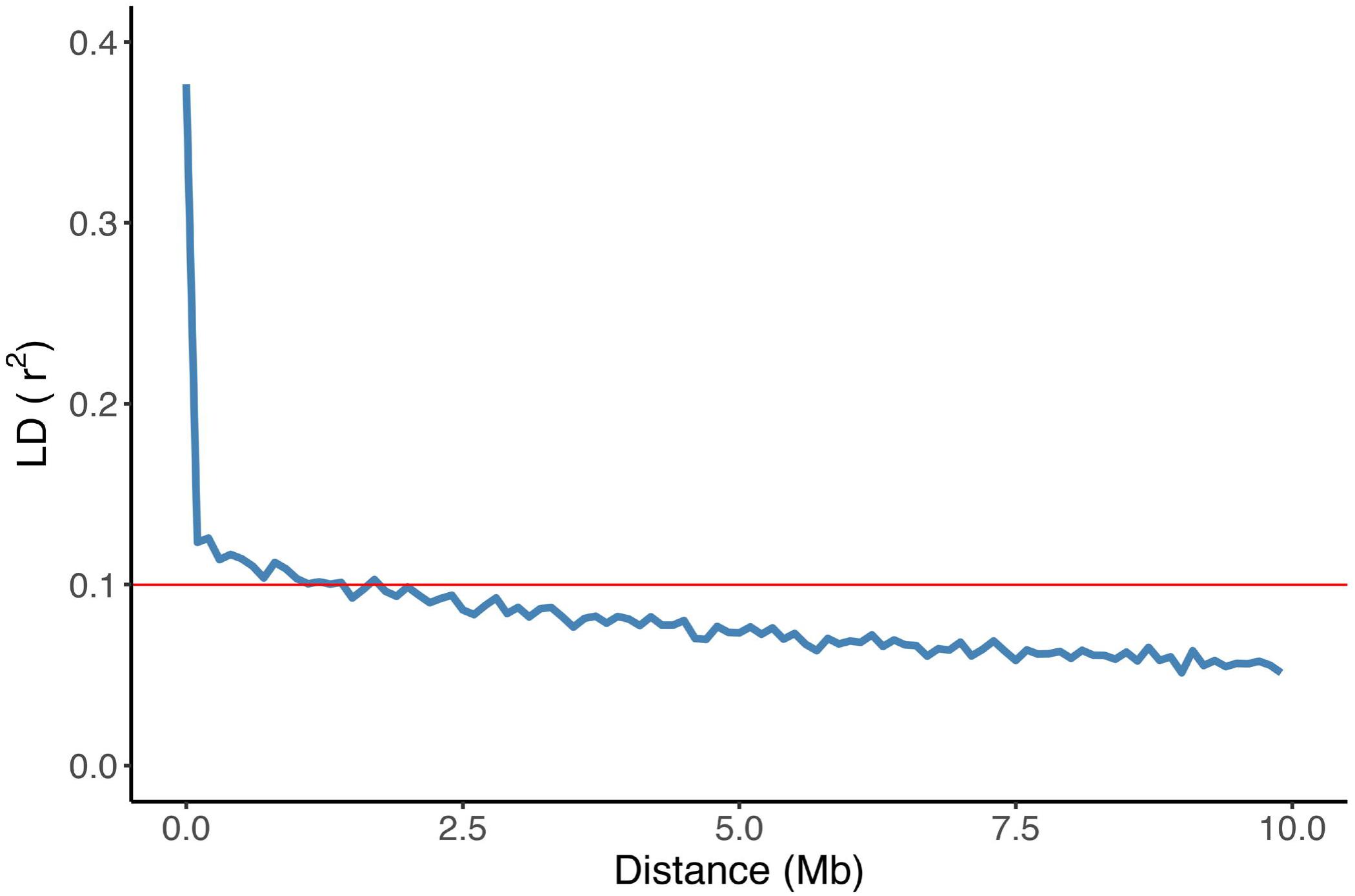
Decay of average LD (r^2^) over distance among SNPs in coho salmon (*Oncorhynchus kisutch*) population. The blue line shows the mean LD in each 1 kb sliding window. The horizontal red line represents significance threshold at 0.1.

**Figure 2.**
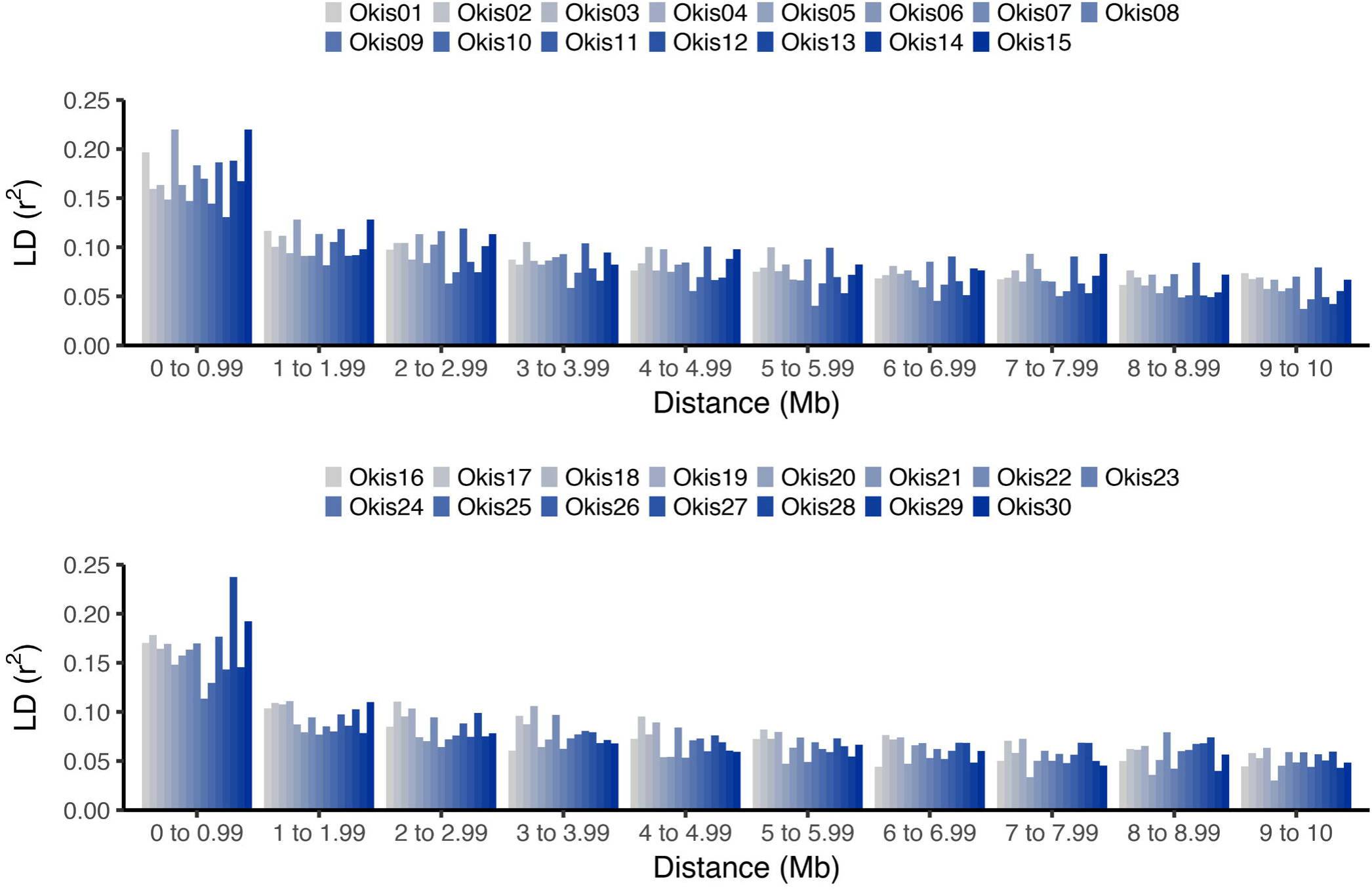
Linkage disequilibrium estimations along the 30 chromosomes of coho salmon. Average values of LD measured as r^2^ per chromosome, according to distances between SNPs. Estimated values are shown from Okis01 to Okis015 (A), and from Okis16 to Okis30 (B)

**Figure 3** illustrates the estimated effective population size of the coho salmon, based on LD, from 8 to 241 generations ago. An increasing Ne as a function of the number of generation was observed, with a Ne of 100 estimated at 10 generations ago, and 1000 for 139 generations ago.

**Figure 3.**
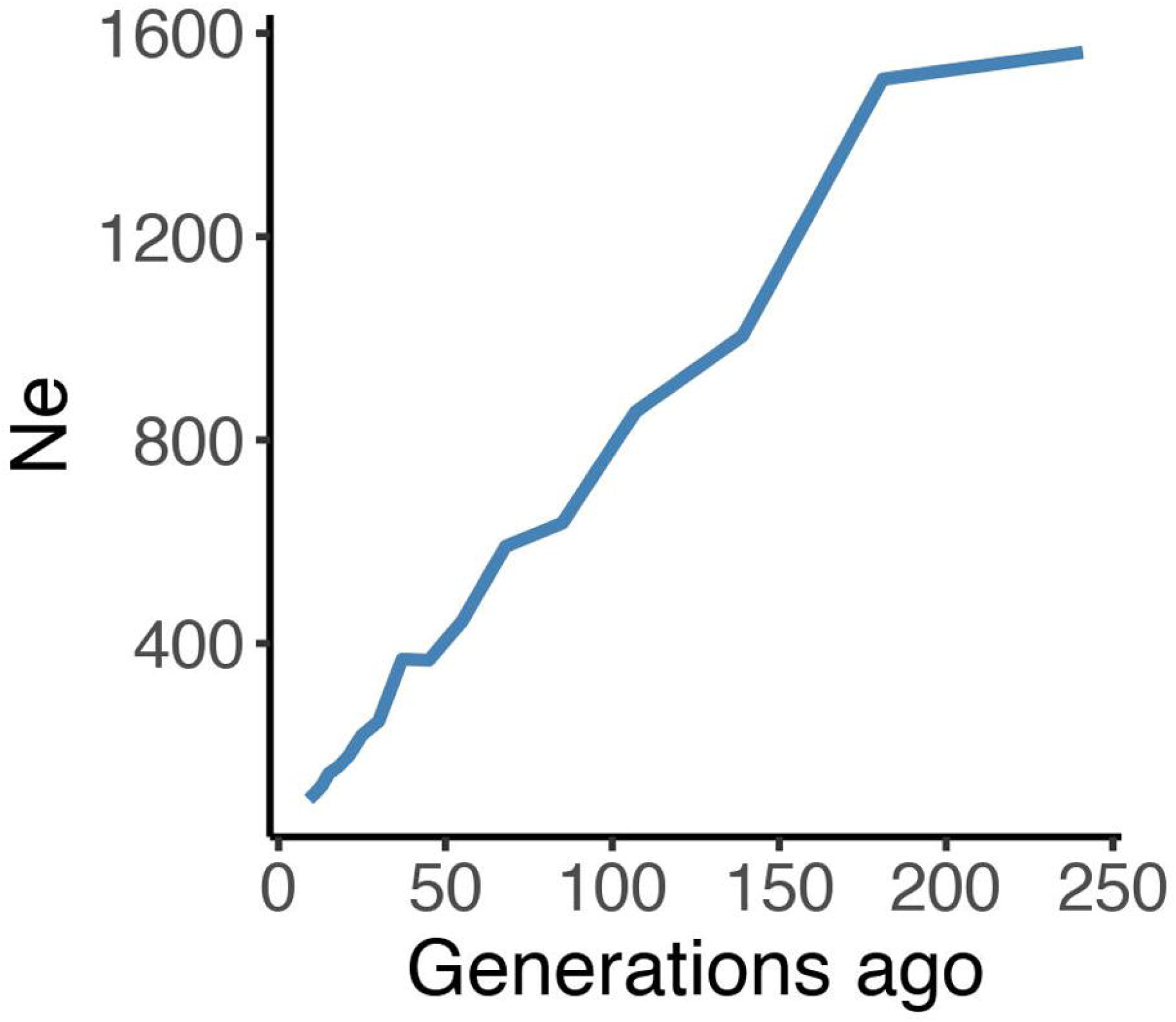
Effective population size estimation in coho salmon population. Estimates of effective population size (Ne) over the past 241 generations based on the LD of an aquaculture strain of coho salmon.

## DISCUSSION

Evaluating the extent and decay pattern of linkage disequilibrium is an important step for statistical genetics. Understanding LD enhances our knowledge of the demographic processes and evolution within the population. Biological factors such as recombination and mutation in conjunction with genetic drift, admixture and effective population size are important variables determining patterns of LD. For this reason, variation in LD among populations and genomic regions are widely reported.

To our knowledge, this is the first study characterizing the LD in a coho salmon population. The samples originated from the broodstock of a breeding program involved with genetically improving coho salmon for production traits. Unrelated animals were chosen in order to avoid LD inflation that can occur with high kinship relationships present (this excluded family-based analyses) (Gutierrez et al., 2015). Due to the increased bias of LD estimations, when estimating |D’| from small sample sizes (Bohmanova et al., 2010), we preferred to use the robust r^2^ statistic. Moreover, to predict the power of association mapping, r^2^ is more useful. The minimum number of individuals necessary for an accurate r^2^ estimation has been suggested to range from 55 to 75 in cattle (Bohmanova et al., 2010; Khatkar et al., 2008). This range increases to 400 or more in case of |D’| (Khatkar et al., 2008). The number of individuals necessary to estimate LD depends on the demographic and genetic population history. Our sample size was within the range suggested above.

Sample sizes above 50 also provide accurate estimations of MAFs (> 0.05) within a population, at a physical distance up to 10 Mb (Khatkar et al., 2008). Filtered markers showed an average MAF of 0.20 per chromosome (**Table 1**). A similar mean value was reported in Nellore cattle, ranging from 0.20 to 0.25 (Espigolan et al., 2013; Matukumalli et al., 2009) and from 0.28 to 0.30 in North American Holstein (Bohmanova et al., 2010). MAFs showed a skewed distribution toward low values, a near identical distribution was found in farmed Tasmanian Atlantic salmon (Kijas et al., 2017). The use of SNP markers with low MAFs tends to underestimate LD measures (Espigolan et al., 2013). LD measurements of r^2^, tend to be less sensitive than |D’| to low MAF (Bohmanova et al., 2010; Khatkar et al., 2008; Kijas et al., 2017).

Estimations of the extent and decay of linkage disequilibrium in the coho salmon breeding population, provide insights into LD patterns in the coho salmon genome, which may have implications for GWAs, GS and for the design of SNP arrays. We estimated the decline in LD, within the population, for values above r^2^ = 0.1 (Delourme et al., 2013; Stich et al., 2013; Vos et al., 2017). For this distance, at least 2,300 SNPs are necessary for whole genome association studies. However, to achieve an accuracy of 0.85 for GEBVs, an average r^2^ greater than 0.2 is required (Meuwissen et al. 2001). At this value, the number of SNPs increases to about 75,000. If we consider a more stringent criterion for higher power genome scans, and consider one SNP every 30 kb, a distance at which the average r^2^ values among SNPs is 0.3 (Ai et al., 2013; Khatkar et al., 2008; Lu et al., 2012), the number of SNPs would increase drastically to 100,000.

Large variation in the average and standard deviation in the LD among chromosomes was found in the current study (**Table 1**). This could be due to variation in recombination rates along different chromosomes (e.g. local hotspots for recombination), decreasing as function of an increase in chromosome length (Arias et al., 2009; Espigolan et al., 2013). Therefore, inferences based on single or only on few chromosomes might be biased and inferences regarding LD would be best when using genome-wide data. LD information from the population may allow researchers to reduce the number of required SNPs for a genomic analysis by excluding redundant SNPs (Khatkar et al., 2008). This can be done by identifying tag SNPs, using information from haplotype block structure, as was previously done in Holstein-Friesian cattle (Khatkar et al., 2007).

Average r^2^ values estimated in our study were higher than those estimated in a wild Finnish Atlantic salmon population, with values ranging from 0.015 to 0.037 (Kijas et al., 2017). However, farmed Tasmanian Atlantic salmon showed mean LD (measured as r^2^) values up to 0.67 for SNPs closer than 1 kb (Kijas et al., 2017), almost double than in the current work (0.37). Linkage disequilibrium estimations in others Atlantic salmon populations, found low LD values, although these estimations were reported in units of recombination (Gutierrez et al., 2015) and using sliding windows of 20 SNPs (Johnston et al., 2014). The different estimation metrics make it difficult to compare directly with the current work. The origin of the current breeding coho population comes from two isolated wild populations (The Kitimat River and Oregon). The admixture of the originated new population may explain the observations of long-range and reduced short-range LD (Pfaff et al., 2001). Pattern that was previously suggested for a Norwegian Atlantic salmon population (Ødegård et al., 2014).

A large decline in Ne was observed ∼ 180 generations ago (approximately 700 years ago, assuming a generation interval of 4 years). This could be due to a significant bottleneck in the wild populations. Similar Ne reductions have been observed in cattle populations (Makina et al., 2015; Villa-Angulo et al., 2009). Even though this is the first study aimed to estimate the effective population size of a coho salmon breeding population, caution must be taken when evaluating the estimations for the number of generations (Corbin et al., 2012). For recent generations, large *c* values are involved and do not necessarily fit the theoretical implications proposed by Hayes (Hayes et al., 2003) for Ne estimations. In the oldest generation after 4Ne generations ago, none of the SNPs can be reliably sampled (Corbin et al., 2012). Therefore, Ne estimations after 4Ne generations ago may be questionable.

## Conclusions

In the current study we used a relatively small sample of coho salmon individuals from a breeding population. We showed the feasibility to estimate LD and infer ancestral population size based on the observed LD using data from ddRAD sequencing. We performed an LD analysis with 62 coho salmon genotyped with 7,505 SNPs. Based on the extent of r^2^ decay of 0.2, we suggest that at least 75,000 SNPs would be necessary for an association mapping study. Increasing this threshold to 0.3, over 100,000 SNPs would be necessary for a high power study, in the current coho salmon population.

## Acknowledgements

AB wants to acknowledge the National Commission of Scientific and Technologic Research (CONICYT) for the funding through the National PhD funding program and to the Government of Canada for the funding through the Canada-Chile Leadership Exchange Scholarship (ELAP).

## Ethics approval and consent to participate

Coho salmon individuals and sampling procedures were approved by the Comité de Bioética Animal from the Facultad de Ciencias Veterinarias y Pecuarias, Universidad de Chile (Certificate N08-2015).

## Authors’ contributions

AB performed DNA extraction, library construction, ddRAD analysis, LD analysis and wrote the initial version of the manuscript. KrC performed library construction and contributed on the data analysis and discussion. GrY contributed with LD analysis and discussion. AJ performed DNA extraction. JPL contributed with study design. WD contributed with analysis and discussion. JMY conceived and designed the study, supervised work of AB and contributed to the analysis, discussion and writing. All authors have reviewed and approved the manuscript.

